# Inhibitors of VPS34 and lipid metabolism suppress SARS-CoV-2 replication

**DOI:** 10.1101/2020.07.18.210211

**Authors:** Jesus A. Silvas, Alexander S. Jureka, Anthony M. Nicolini, Stacie A. Chvatal, Christopher F. Basler

## Abstract

Therapeutics targeting replication of SARS coronavirus 2 (SARS-CoV-2) are urgently needed. Coronaviruses rely on host membranes for entry, establishment of replication centers and egress. Compounds targeting cellular membrane biology and lipid biosynthetic pathways have previously shown promise as antivirals and are actively being pursued as treatments for other conditions. Here, we tested small molecule inhibitors that target membrane dynamics or lipid metabolism. Included were inhibitors of the PI3 kinase VPS34, which functions in autophagy, endocytosis and other processes; Orlistat, an inhibitor of lipases and fatty acid synthetase, is approved by the FDA as a treatment for obesity; and Triacsin C which inhibits long chain fatty acyl-CoA synthetases. VPS34 inhibitors, Orlistat and Triacsin C inhibited virus growth in Vero E6 cells and in the human airway epithelial cell line Calu-3, acting at a post-entry step in the virus replication cycle. Of these the VPS34 inhibitors exhibit the most potent activity.

## INTRODUCTION

SARS-CoV-2, a member of the *Betacoronavirus* genus, is an enveloped positive-sense, RNA virus responsible for a current pandemic^1^. Because of its profound impact on society and human health there is an urgent need to understand SARS-CoV-2 replication requirements and to identify therapeutic strategies^2^. Repurposing drugs developed for other purposes may provide a shortcut to therapeutic development^3–6^. The use of compounds known to target specific host factors may also elucidate key pathways needed for virus replication.

Coronavirus (CoV) replication involves multiple critical interactions with host cell membranes, including during viral entry and virus release^2, 7–9^. In addition, one of the most striking features of CoV infection is the establishment of replication organelles that consist of double membrane vesicles (DMV), double-membrane spherules (DMSs) and convoluted membranes (CM) with DMVs serving as the main site of viral RNA synthesis^10^. The origin of these membrane organelles in beta-coronavirus infection remains incompletely understood. The membrane structures colocalize with LC3, a protein with well-known functions in autophagy^7, 11^. In murine embryonic stem cell lines, autophagy was found to be critical for DMV formation and replication of the beta-coronavirus mouse hepatitis virus^7^. However, studies in bone marrow derived macrophages or primary mouse embryonic fibroblasts lacking ATG5 indicated that autophagy is not essential for DMV formation or MHV replication^11^. An alternate model indicates that beta coronaviruses usurp vesicles known as EDEMosomes, which associate with non-lipidated LC3 and normally function to regulate ER-associated degradation (ERAD), to provide membranes for replication^8^.

Many enveloped, positive-sense RNA viruses that replicate in double membrane compartments have been demonstrated to be sensitive to inhibitors of various aspects of membrane metabolism/biology. For example, VPS34 a class III phosphoinositol-3 kinase (PI3K) that plays roles in autophagy, endosomal trafficking, and other aspects of membrane biology has been implicated in the replication of hepatitis C virus (HCV) and tombusvirus (TBSV)^12, 13^. The compound Triacsin C, which inhibits an enzyme upstream of triglyceride synthesis, long chain fatty acyl CoA, impairs the growth of several viruses that require for replication lipid droplets, organelles that serve as storage sites for neutral lipids such as triacylglycerol^14–16^. Downstream of long chain fatty acyl CoA in the synthesis of triglycerides are diacylglycerol acyltransferases 1 and 2 (DGAT1 and DGAT2). Inhibition of these enzymes inhibits HCV and rotavirus replication. More general inhibitors of fatty acid synthetase such as Orlistat, also decrease replication of several different viruses^17–20^.

Here we asked whether SARS-CoV-2 is susceptible to modulators of lipid metabolism by assessing the sensitivity of the virus in Vero E6 and Calu-3 cells to VPS34 inhibitors, Triacsin C, inhibitors of DGATs and Orlistat, an inhibitor of FASN^21^. We find that two inhibitors of VPS34 potently inhibited SARS-CoV-2 replication, whereas an FDA-approved inhibitor of a different class of PI3K had minimal effect on replication. Targeting FASN and *de novo* synthesis of triacylglycerol, diacylglycerol and cholesterol esters each impairs SARS-CoV-2 replication whereas inhibition of DGATs was not effective. We also identified that each inhibitor exhibits antiviral effects post-entry and that they perturb the structure of viral replication centers. Taken together, the data presented here implicates specific lipid metabolism pathways in SARS-CoV-2 replication and suggests that these pathways are promising therapeutic targets.

## MATERIALS AND METHODS

### Virus and cell lines

Vero E6 (ATCC# CRL-1586), Calu-3 (ATCC# HTB-55), and Caco-2 (ATCC# HTB-37) were maintained in DMEM (Corning) supplemented with 10% heat inactivated fetal bovine serum (FBS; GIBCO). Cells were kept in a 37°C, 5% CO_2_ incubator without antibiotics or antimycotics. SARS-CoV-2, strain USA_WA1/2020, was obtained from the World Reference Collection for Emerging Viruses and Arboviruses at the University of Texas Medical Branch-Galveston.

### Virus Propagation and Plaque Assays

A lyophilized ampule of SARS-CoV-2 was initially resuspended in DMEM supplemented with 2% FBS. VeroE6 cells were inoculated in duplicate with a dilution of 1:100 with an adsorption period of 1 hour at 37C and shaking every 15 minutes. Cells were observed for cytopathic effect (CPE) every 24 hours. Stock SARS-CoV-2 virus was harvested at 72 hours post infection (h.p.i) and supernatants were collected, clarified, aliquoted, and stored at −80°C.

For plaque assays, Vero E6 cells were seeded onto a 24-well plate 24 hours before infection. 100ul of SARS-CoV-2 serial dilutions were added, adsorbed for 1 hour at 37C with shaking at 15-minute intervals. After the absorption period, 1 mL of 0.6% microcrystalline cellulose (MCC; Sigma 435244) in serum-free DMEM was added. To stain plaque assays MCC was removed by aspiration, and 10% neutral buffered formalin (NBF) added for one hour at room temp and then removed. Monolayers were then washed with water and stained with 0.4% crystal violet. Plaques were quantified and recorded as plaque forming units (PFU)/mL.

### Confocal microscopy

For confocal microscopy analysis, all cell lines were pre-seeded 24 hours before infection onto glass coverslips and infected with SARS-CoV-2 at a multiplicity of infection (MOI) of 1. At 24 hours post-infection (h.p.i.) supernatant was removed, and samples fixed with 10% NBF for 1 hour at room temperature followed by PBS wash and permeabilized with sterile filtered 0.1% Saponin in PBS. Cells were blocked with 0.1% Saponin in Fluorescent Blocker (ThermoFisher) for 1 hour at RT. Primary antibodies were added and incubated overnight at 4C. AlexaFluor488, 594, and 647 conjugated secondary antibodies were used and nuclei stained with DAPI. Samples were imaged on Zeiss LSM800 Confocal with Super Resolution AiryScan. Images were rendered in ZenBlue or Imaris Viewer 9.0.

### Maestro Z Impedance Experiments

Prior to cell plating, CytoView-Z 96-well electrode plates (Axion BioSystems, Atlanta, GA) were coated with 5 μg/mL human fibronectin (Corning) for 1 hr at 37C. After coating, fibronectin was removed and 100 μL of DMEM/10% FBS was added to each well. The plate was then docked into the Maestro Z instrument to measure impedance electrode baseline. Vero E6 cells were then plated to confluency (~75,000 cells/well) in the coated CytoView-Z plates and left at room temperature for 1 hour to ensure even coverage of the well. Plates containing Vero E6 cells were then docked into the Maestro Z for 24 hours at 37°C/5% CO_2_ to allow the cells to attach and the monolayer to stabilize, as measured by resistance, a component of impedance. The Maestro Z was used to monitor the resistance of the monolayer as it formed, very similar to transepithelial electrical resistance (TEER)^22^. In this study, resistance was measured at 10 kHz, which reflects both cell coverage over the electrode and strength of the barrier formed by the cell monolayer. For compound treatments, media was removed from wells of the CytoView-Z plates and 195 μL of pre-warmed DMEM/2% FBS was added with the indicated concentration of compound. Infections with SARS-CoV-2 at an MOI of 0.01 were carried out by directly adding 5 μL of virus to each well. Plates were then docked within the Maestro Z and resistance measurements were continuously recorded for 48-72 hours post-infection. All plates contained media only, full lysis, uninfected, and SARS-CoV-2 infected controls. For calculation of percent inhibition, raw resistance values for each well were normalized to the value at 1 hour post-infection within the Axis Z software, and percent inhibition was calculated with the following formula: Percent Inhibition = 100*(1-(1-average of treated cells)/(1-average of infected control)). Median time to death calculations were performed by fitting the Boltzmann sigmoid equation to raw kinetic resistance data in Graphpad Prism. Fifty percent maximum velocity (V50) values obtained from the Boltzmann sigmoid fits were used to determine median time to death for each MOI.

### Cell viability assay

VeroE6 or Calu-3 cells were seeded in 96-well black walled microplates and incubated overnight. Cells were then treated with compounds and CellTox Green Dye (Promega) to monitor compound cytotoxicity. Fluorescence (Excitation: 485nm, Emission: 520nm) was measured every 24 hours post treatment for 3 days. Percent viability was determined using the minimum fluorescence obtained from media only cells and the maximum value obtained by cells lysed with 1% Triton-X.

### Labeling of nascent viral RNA

VeroE6 cells were seeded onto glass coverslips and incubated overnight at 37C. Cells were then infected with SARS-CoV-2 at an MOI of 3. At 24 h.p.i. cells were treated with 1μM of Actinomycin D (Sigma) for 1 hour. Nascent RNA was labeled using Click-iT™ RNA Alexa Fluor™ 594 Imaging Kit (ThermoFisher). Cells were then processed for confocal analysis.

### Compounds

VPS34 IN-1 (#17392), PIK-III (#17002), Triacsin C (#10007448), and Orlistat (#10005426) were purchased from Cayman Chemical (Ann Arbor, Michigan). Remdesivir was purchased from Target Molecule Corp. (T7766, Boston, Massachusetts). T863 (#SML0539) and PF06424439 (#PZ0233) were purchased from Sigma-Aldrich (St. Louis, Missouri). All chemicals were resuspended in dimethylsulfoxide (DMSO). ^23^

## RESULTS

### Development of 96-well format assay to measure SARS-CoV-2 cytopathic effects

SARS-CoV-2 induces significant cytopathic effects in infected Vero E6 cells. Based on this property, we standardized a 96-well format assay that provides continuous real-time, label-free monitoring of the integrity of cell monolayers, thereby providing assessment of virus growth through decreased cell viability. This assay was standardized using the Maestro Z platform (Axion BioSystems, Atlanta, GA), an instrument that uses 96-well plates containing electrodes in each well (CytoView-Z plates). The electrodes measure electrical impedance across the cell monolayer every minute throughout the course of the experiment. As SARS-CoV-2 replication damages the cell monolayer, impedance measurements decrease over time, providing a detailed assessment of infection kinetics.

The capacity of the system to differentiate different levels of virus replication was first assessed. Confluent Vero E6 monolayers in CytoView-Z plates were infected with SARS-CoV-2 at multiple MOIs (10 to 0.0001) and resistance measurements were acquired for 72 hours post-infection. As shown in **Figure 1A**, the progression of infection at each MOI was clearly distinct. A decrease in resistance could be observed as early as 18-20 h.p.i. at an MOI of 10 and 1, and as late as 56 h.p.i. at an MOI of 0.0001. Depending on MOI, signals reached their nadirs between 32 to 72 h.p.i. To correlate with a decrease in resistance, the raw kinetic data was used to determine the median time to cell death for each MOI **(Figure 1B)**. Based on its desirable kinetics, the MOI of 0.01 was chosen for the screening of compounds for antiviral activities.

**Figure 1.**
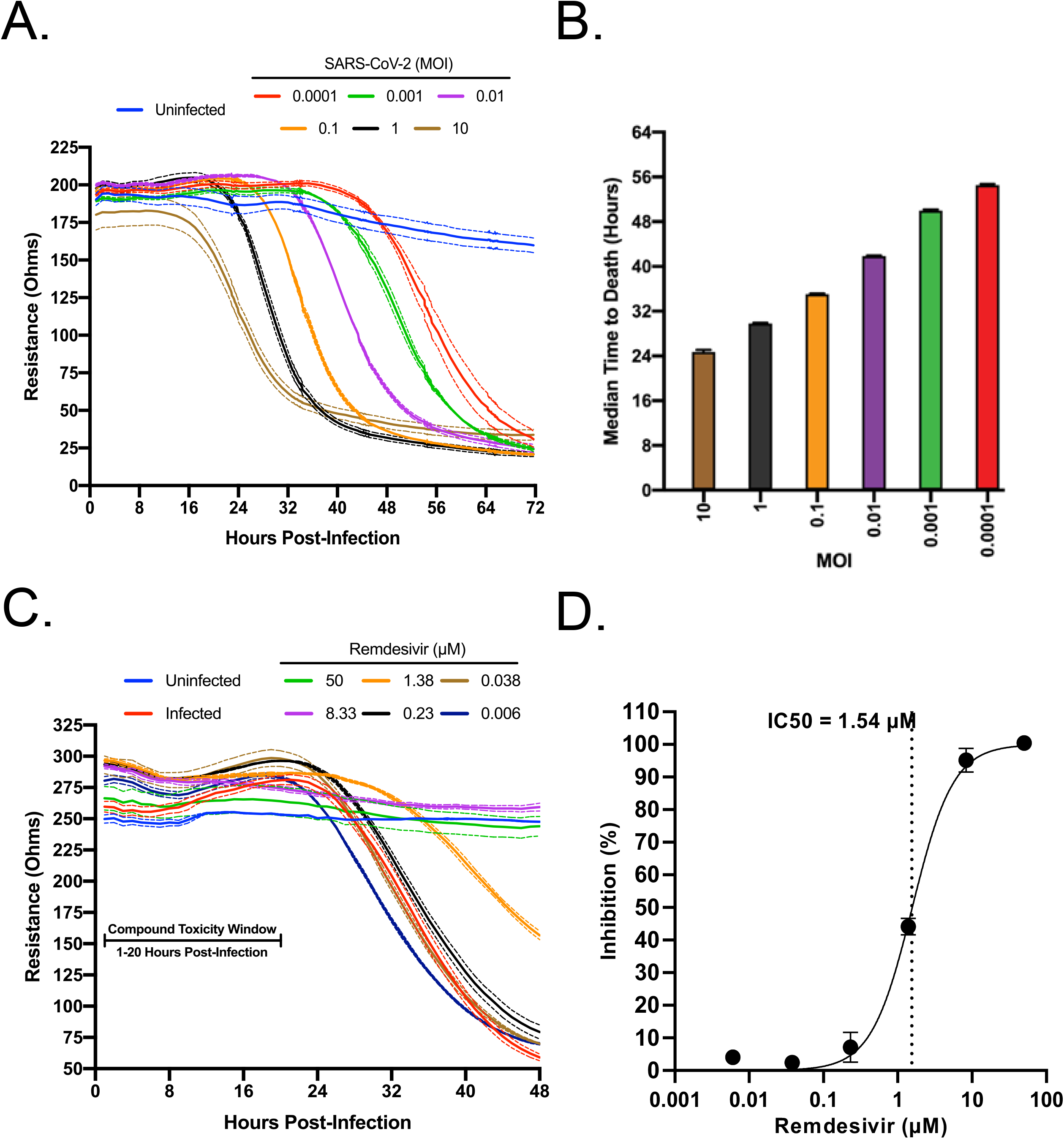
Standardization of an electrical resistance-based assay as a measure of SARS-CoV-2 induced CPE and anti-SARS-CoV-2 activity. VeroE6 cells were seeded into a CytoView-Z 96-well plate and cells were allowed to stabilize overnight, as measured by electrical resistance. **A)** SARS-CoV-2 was titrated in 10-fold dilutions ranging from 10-0.0001 MOI. Resistance was measured every minute over the course of 72 hours. Solid lines indicate the mean, dotted lines indicate the standard error of three replicates. **B)** Median time to death calculations based on raw resistance data for each MOI. **C)** Remdesivir was titrated in 6-fold dilutions ranging from 50-0.006 μM. After infection at an MOI of 0.01, resistance was monitored for 48 h.p.i. and **D)** percent inhibition was determined at the 48 hour timepoint.

To establish the Maestro Z as a potential instrument for screening of anti-SARS-CoV-2 therapeutics, we first tested Remdesivir, a well-described inhibitor of SARS-CoV-2 that has been granted emergency use authorization (EUA) for the treatment of COVID-19^24, 25^. Vero E6 cells were seeded on a CytoView-Z plate, incubated overnight to allow cells to stabilize, pretreated with 6-fold dilutions of Remdesivir for 1 hour and infected with SARS-CoV-2. Resistance measurements were recorded for 48 h.p.i. **(Figure 1C)**. In agreement with previous studies, we determined an 50% inhibitory concentration (IC50) for Remdesivir of 1.54 μM **(Figure 1D)**^24^. Taken together, these data validate the impedance-based assay described as a tool for screening of potential SARS-CoV-2 therapeutics.

### Inhibitors of VPS34 activity impair SARS-CoV-2 growth

VPS34 is a multifunctional protein involved in autophagy and membrane trafficking. Since coronaviruses induce formation of double membrane vesicles for replication, we wanted to determine if VPS34 activity was essential for SARS-CoV-2 replication. Therefore, we tested two well characterized VPS34 inhibitors IN-1 (referred as VPS34-IN1 below) and PIK-III over a 10-point dose response in the resistance assay^26^. The compounds were added to pre-plated Vero E6 cells 1 hour prior to infection with SARS-CoV-2 at a MOI of 0.01. Both VPS34-IN1 and PIK-III induced rapid cytotoxicity at 50 μM and 16.67 μM as indicated by a rapid decrease in resistance measurements between 1 and 20 h.p.i. **(Figure 2A and 2C)**. However, at concentrations of 5.56 μM and below, the integrity of the monolayer was preserved relative to the mock-treated control indicating an antiviral effect and an absence of cytotoxicity. Calculations based on normalized resistance measurements at 48 h.p.i for non-toxic doses yielded IC50s of 0.29uM for VPS34-IN1 and 0.202uM for PIK-III **(Figure 2B and 2D, respectively)**. Additionally, IC90s of 2.52 μM (VPS34-IN1) and 1.81 μM (PIK-III) were also calculated. These data suggest that the VPS34 kinase plays a significant role in SARS-CoV-2 replication and is a potential target for therapeutic intervention.

**Figure 2.**
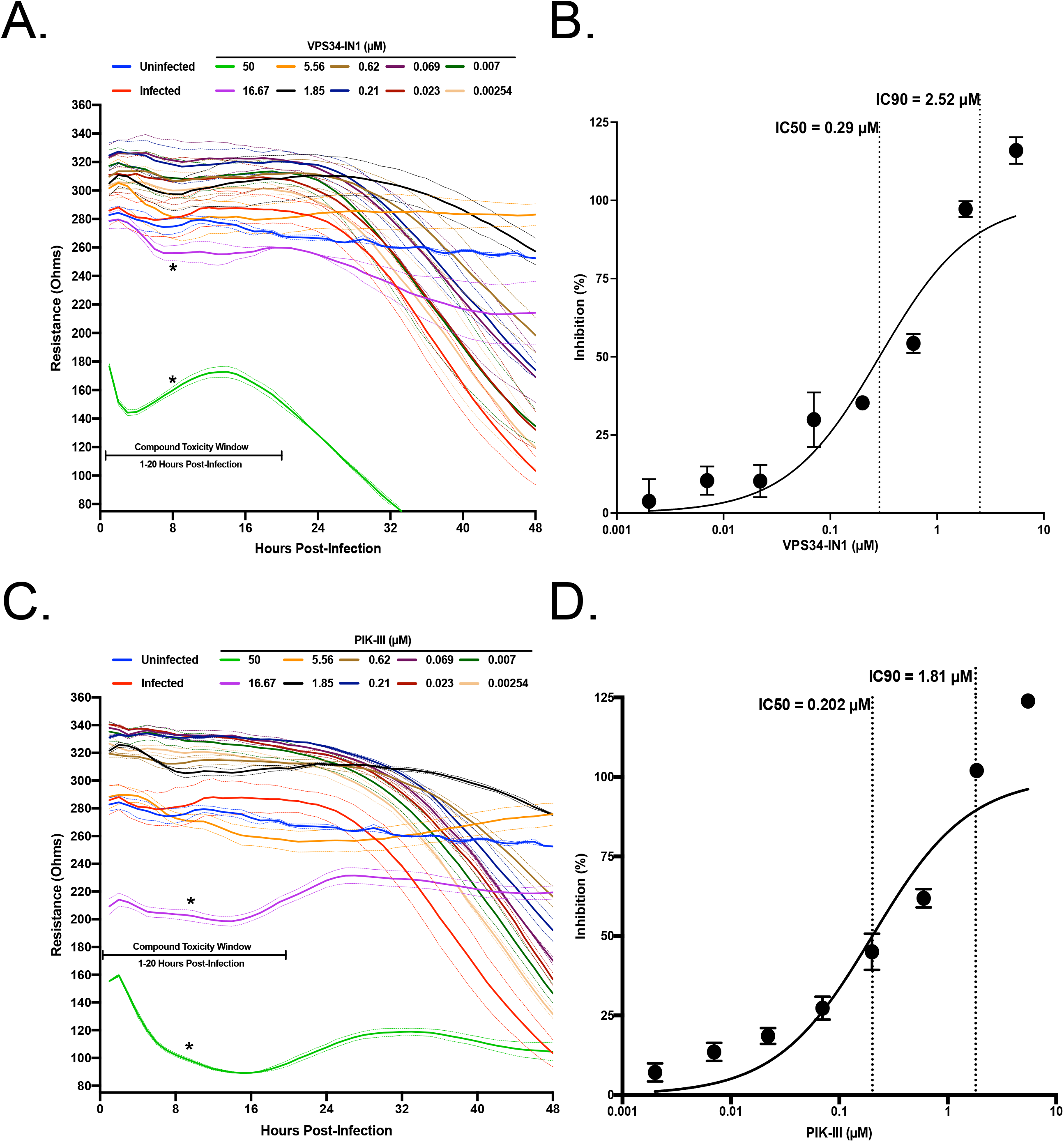
VPS34 inhibitors exhibit anti-SARS-CoV-2 activity. VeroE6 cells were seeded into a CytoView-Z 96-well plate, and cells were allowed to stabilize overnight. Cells were pre-treated with serial half-log dilutions of **A)** VPS34-IN1 or **C)** PIK-III and infected with SARS-CoV-2 at an MOI=0.01. Resistance **(A and C)** was measured every minute over the course of 48 hours and percent inhibition **(B and D)** was determined at the 48-hour timepoint. Solid lines indicate mean, dotted lines indicate the standard error of two replicates.

### Inhibition of fatty acid metabolism inhibits SARS-CoV-2 replication

Fatty acid metabolism leads to production of triglycerides, phospholipids and other molecules^27^. Elongation of the phospholipid membranes can be aided by channeling fatty acid into phospholipid synthesis^28^. Modulation of fatty acid metabolism has been shown to impact several viruses such as dengue virus, hepatitis C virus, and Old World alphaviruses^18, 29, 30^. Two well-described compounds that inhibit fatty acid metabolism are Orlistat and Triacsin C, both of which have been shown to have antiviral activity^19, 30^. Orlistat is an FDA-approved drug that inhibits lipases and also fatty acid synthase (FASN), and Triacsin C inhibits long chain Acyl-CoA synthetases. To test these against SARS-CoV-2, VeroE6 cells were pre-seeded onto a CytoView-Z plate, allowed to stabilize and then pre-treated with Triacsin C or Orlistat for 1 hour before infection with SARS-CoV-2 at an MOI of 0.01. Based on the toxicity window of 1-20 h.p.t. determined with the VPS34 inhibitors, neither Triacsin C nor Orlistat induced early cytotoxic effects, even at the highest concentrations of 50uM and 500uM, respectively **(Figure 3A and 3C)**. Both compounds exhibited inhibition at the higher concentrations tested, although complete inhibition was not achieved even with 500 μM of Orlistat. Based on the data we extrapolated an IC50 of 422.3uM for Orlistat and calculated an IC50 of 19.5uM for Triacsin C **(Figure 3B and 3D)**. Viruses such as HCV and rotavirus that are sensitive to inhibition by Triacsin C are also impaired by inhibitors of DGATs^14, 31^. Therefore, we tested the effects of DGAT1 and DGAT2 inhibitors T863 and PF06424439^32, 33^. Neither compound displayed any inhibitory activity **(Supplemental Figure 1)**. This data suggests that metabolism of fatty acids plays an important role in SARS-CoV-2 infection.

**Figure 3.**
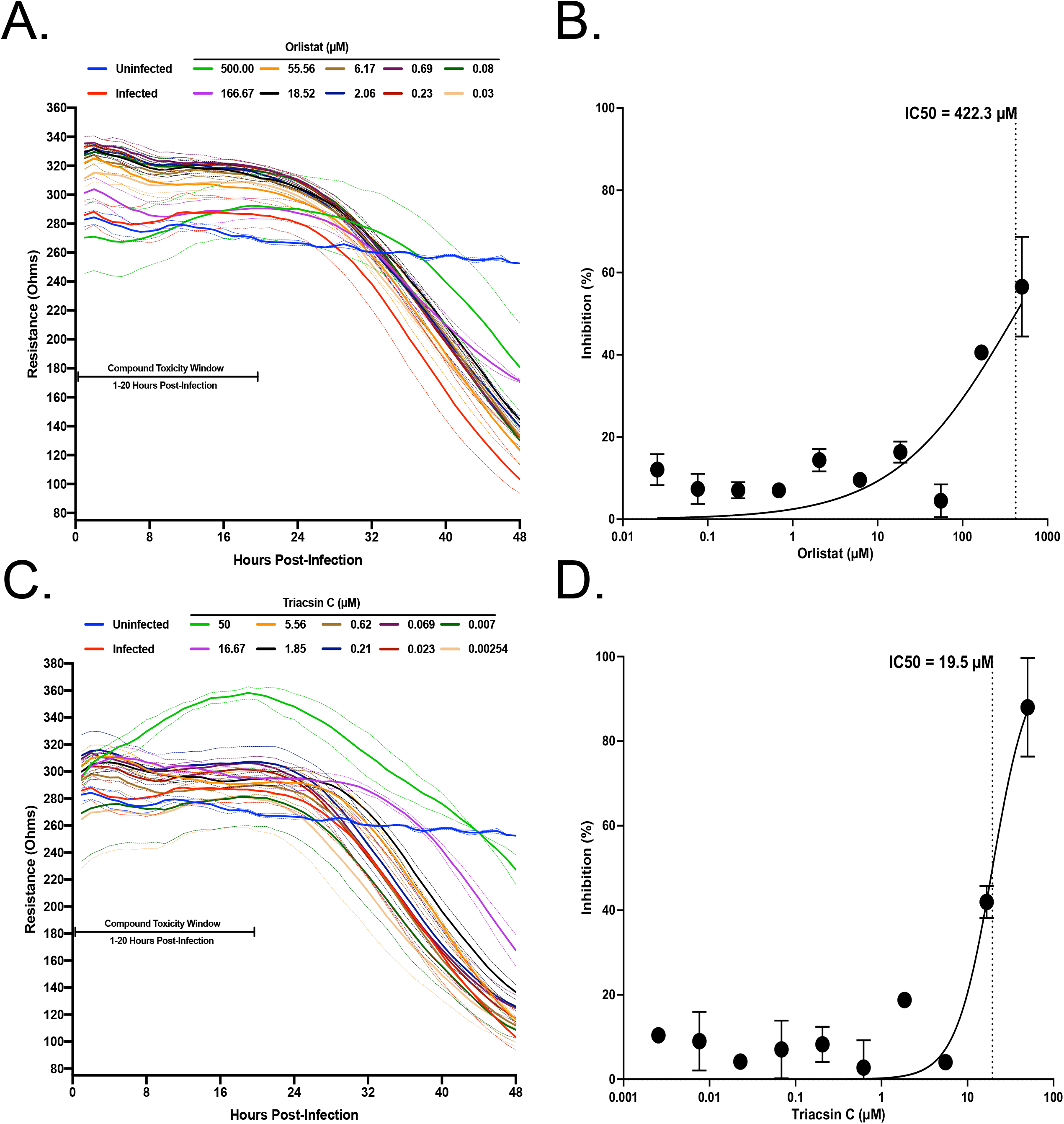
Screening of fatty acid inhibitors for potential anti-SARS-CoV-2 activity. VeroE6 cells were seeded into a CytoView-Z 96-well plate and allowed to stabilize overnight. Cells were pre-treated with serial half-log dilutions of **A)** Orlistat or **B)** Triacsin C and infected with SARS-CoV-2 at an MOI=0.01. Resistance **(A and C)** was measured every minute over the course of 48 hours and percent inhibition **(B and D)** was determined at the 48-hour timepoint. Solid lines indicate the mean and dotted lines indicate the standard error of two replicates.

### VPS34 inhibitors exhibit potent attenuation of SARS-CoV-2 early and late in its replication cycle

Next, time-of-addition studies were performed. We sought to determine how long the addition of VPS34-IN1, PIK-III, Orlistat, or Triacsin C could be postponed before activity was lost. Additionally, this would identify if the anti-viral activity of each compound impacted a pre- or post- viral entry step. As indicated in **Figure 4A**, 4 conditions were tested 1) single treatment 1 hour prior to viral infection, with compound removed just prior to infection; 2) 1 hour pre-treatment with continuous dosing; 3) dosing at 2 h.p.i.; and 4) dosing at 4 h.p.i.. VeroE6 cells were pre-seeded onto a CytoView-Z plate and allowed to stabilize, compounds were added, and resistance was monitored for 48 hours after infection. Percent inhibition was calculated based on resistance values at 48 h.p.i. We observed that a single 5 μM treatment of VPS34-IN1 or PIK-III inhibited SARS-CoV-2 replication **(Figure 4B)**. Additionally, inhibition was observed even when added after 4 h.p.i. In contrast, removal of Orlistat or Triacsin C before infection, eliminated their efficacy. Maintenance throughout the experiment was inhibitory, as was addition at 2 or 4 hours post infection. Interestingly, delayed treatment with Triacsin C at 50μM exhibited greater anti-viral activity that initiating the treatment one hour prior to infection. Altogether, these data demonstrate activity of the VPS34 inhibitors at both early and late, post-entry time points and indicate that the effects of Orlistat and Triacsin C are likely post-entry.

**Figure 4.**
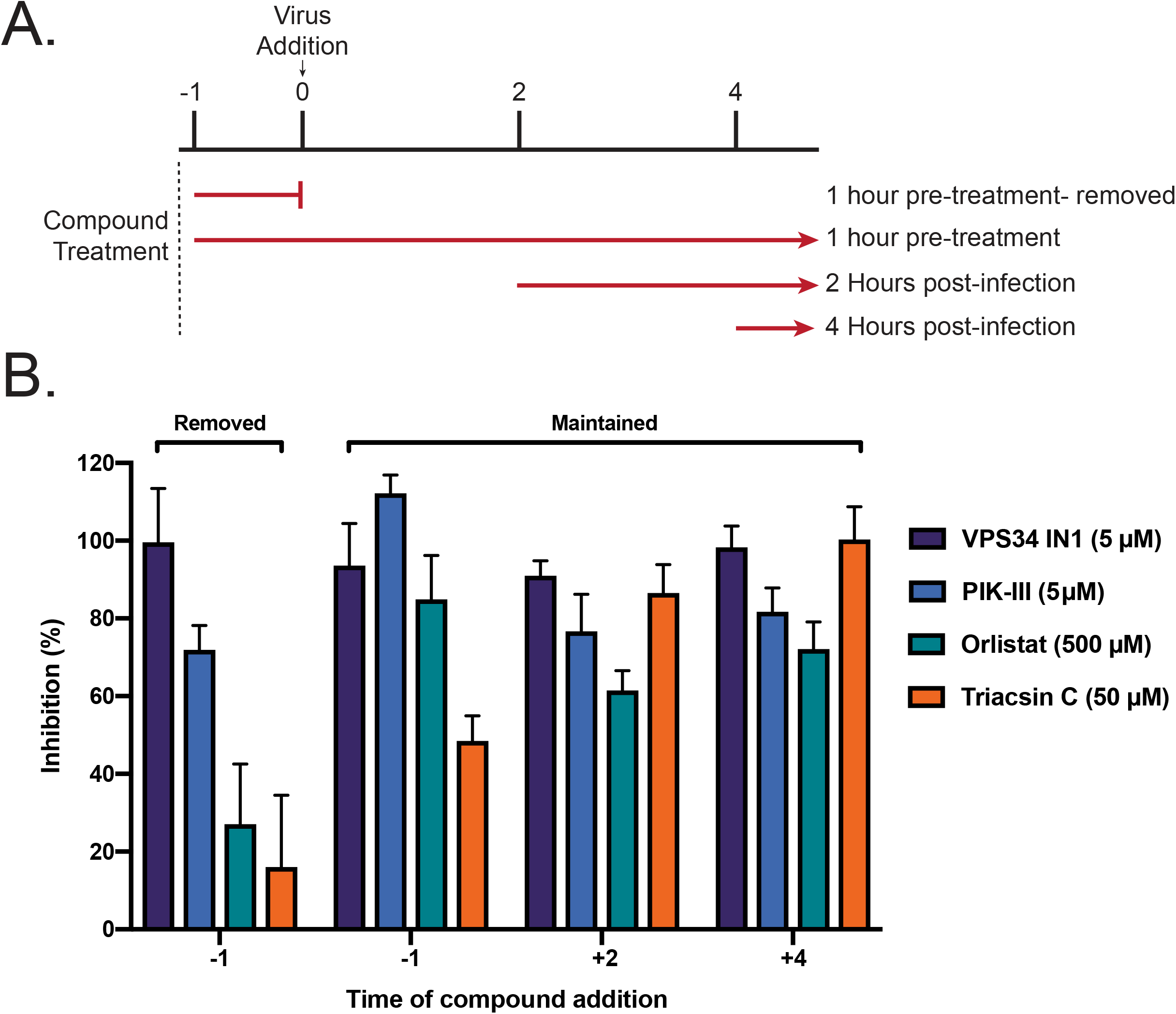
Single treatment of VPS34 inhibitors have potent anti-viral activity against SARS-CoV-2. VeroE6 cells were seeded into a CytoView-Z 96-well plate, and allowed to stabilize overnight. **A)** Timeline for the time-of-addition experiment. **B)** VeroE6 cells were pre-treated for one hour and compound was removed (−1), pre-treated for one hour with compound maintained throughout infection (+1), or treated at 2 (+2) or 4 (+4) hours post-infection with an MOI of 0.01. Resistance was measured every minute over the course of 48 hours and percent inhibition was determined at the 48-hour timepoint. Data is representative of the mean and standard error of three technical replicates.

### Attenuation of VPS34 kinase activity and fatty acid metabolism inhibit SARS-CoV-2 in a human airway epithelial cell line

We proceeded to investigate if the inhibitors were effective in the human lung carcinoma cell line, Calu-3, by directly measuring production of infectious virus and cytotoxicity. That this cell line is derived from the human airway and is highly susceptible to infection has established it as a standard for infection studies with SARS-CoV-1, MERS-CoV and SARS-CoV-2^34, 35^. Calu-3 cells were plated onto 96-well plates and allowed to reach 95% confluency. Cells were then pre-treated with a range of concentrations of VPS34-IN1, PIK-III, Triacsin C, Orlistat, DMSO, or mock treated with media alone for 1 hour then infected with SARS-CoV-2 at an MOI of 0.01. Supernatants were collected at 48 h.p.i. and titered on VeroE6 cells by plaque assay. In parallel, to determine cytotoxicity of these compounds, Calu-3 cells were seeded onto 96-well black walled 96-well plates, allowed to reach 95% confluency and treated with VPS34-IN1, PIK-III, Triacsin C, Orlistat, DMSO, or mock treated with media alone. CellTox Green was added at the time of dosing and fluorescence measured at 48 h.p.i. in order to assess cytotoxicity. Each of the compounds inhibited production of infectious virus, as measured by plaque assay on Vero E6 cells **Figure 5A, C, E, and G)**. In contrast to VeroE6 cells, no cytotoxicity was observed even at the highest dose for each compound in Calu-3 cells. We observed IC50s of 0.55μM (VPS34-IN1), 0.12μM (PIK-III), 21.25μM (Orlistat), and 0.04μM (Triacsin C), as shown in **Figure 5B, D, F, and H**, respectively. Importantly, the IC50s calculated for VPS34-IN1 and PIK-III by measuring infectious virus are in close agreement with IC50s calculated in Vero E6 cells using the resistance-based assay. The IC50s for Triacsin C and Orlistat were substantially lower than in the Vero cells. These data suggest that attenuation of the kinase activity of VPS34, synthesis of fatty acids or production of long chain fatty acyl-CoA in human bronchial epithelial cells inhibits replication of SARS-CoV-2.

**Figure 5.**
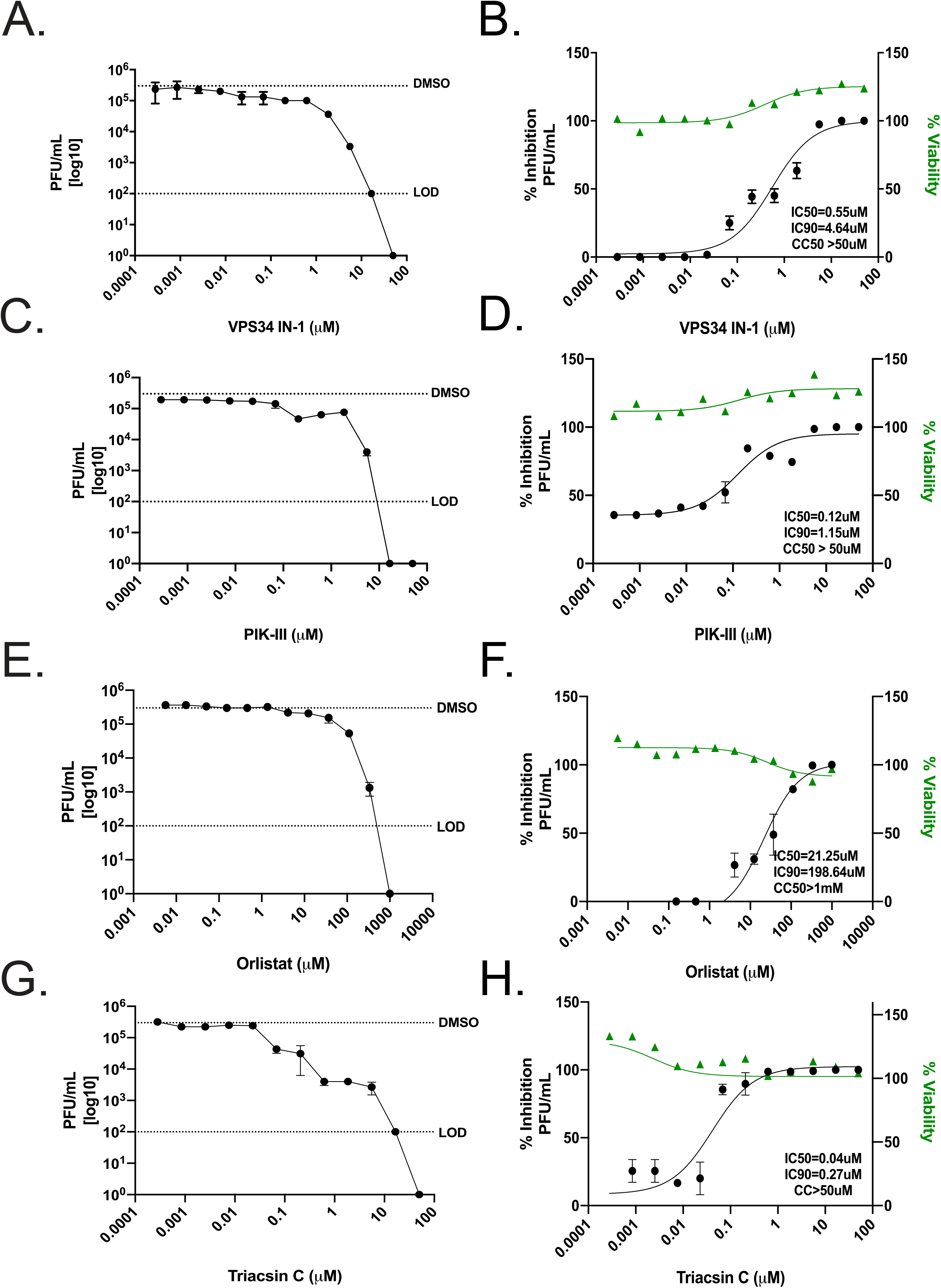
Attenuation of VPS34 kinase activity and fatty acid metabolism inhibit SARS-CoV-2 replication in human airway epithelial cell line. Calu-3 cells were plated onto a 96-well plate and allowed to reach 95% confluency. Cells were then pre-treated with a range of concentrations of **A-B)** VPS34-IN1, **C-D)** PIK-III, **E-F)** Orlistat, **G-H)**, Triacsin C, DMSO, or mock-treated with media alone for 1 hour then infected with SARS-CoV-2 at an MOI of 0.01. Supernatants were collected at 48 h.p.i. and virus was quantified by plaque assay on VeroE6 cells. The data is reported as plaque forming units per milliliter (pfu/ml) **(left panels)**. Cell viability over 48 hours was determined in parallel. Percent inhibition, IC50, and IC90 were calculated from the plaque assay data and plotted with the cell viability data **(right panels)**. The dotted line labeled DMSO indicates the level of virus growth in the DMSO control. The dotted line labeled LOD indicates the limit of detection of the plaque assay.

VPS34 is a class III PI3 kinase. We therefore extended our study to determine if BYL719, an FDA approved inhibitor of class I PI3 kinase used to treat breast cancer, would also inhibit SARS-CoV-2 replication in Calu-3 cells. Unlike the VPS34-specific inhibitors, little inhibition was detected up to 16.6 μM, at which we observed a 1-log decrease in viral titers **(Supplemental Figure 2)**. This data suggests that not all PI3K classes play a significant role during SARS-CoV-2 replication.

### Inhibition of VPS34 kinase activity and fatty acid metabolism disperse SARS-CoV-2 replication centers

SARS-CoV-1 and MERS-CoV replicate in double membrane compartments to which the autophagy membrane marker LC3 localizes^8, 9, 23^. We investigated if, similar to SARS-CoV-1 and MERS, SARS-CoV-2 nascent viral RNA and N co-localized with LC3. VeroE6 cells were infected with SARS-CoV-2 at a MOI of 3 and at 24 h.p.i., were treated with 1μM of actinomycin D to arrest host-cell transcription. Cells where then chased for 4 hours with 5-ethynyl uridine (EU). Viral nascent RNA labeled during the EU chase was then detected with click chemistry, indirect immunofluorescence performed using primary antibodies against N and LC3, and the endoplasmic reticulum (ER) was detected with DPX BlueWhite ER stain. We observed distinct formation of ring-like structures positive for ER, N, LC3, and nascent viral RNA **(Supplemental Figure 3A)**. Co-localization analysis demonstrated that nascent viral RNA co-localized with N or LC3 **(Supplemental Figure 3B)**. This data demonstrates the presence of SARS-CoV-2 replication centers that form in association with LC3.

Because each compound exhibited inhibitory effects when added after viral entry, we next asked whether the compounds altered the establishment of viral replication centers. Calu-3 cells were seeded onto fibronectin coated glass cover slips and allowed to reach 95% confluency. Cells were pre-treated with approximately the IC90 of VPS34-IN1 (5 μM), PIK-III (5 μM), Orlistat (500 μM), or Triacsin C (50 μM) and infected with SARS-CoV-2 at a MOI of 3. At 24 h.p.i. cells were fixed, permeabilized, and indirect immunofluorescence performed using primary antibodies against SARS-CoV-2 nucleoprotein (N) and dsRNA. We observed that when compared to the media only or DMSO controls, N became completely cytoplasmic and did not form any large inclusion like formations in the presence of the compounds **(Figure 6)**. Additionally, even though dsRNA could be detected both distributed throughout the cytoplasm and associated with N in large inclusion like formations in the media only and DMSO controls, in the cells treated with inhibitors, dsRNA was only found distributed throughout the cytoplasm. This data suggests that the compound disrupt replication center formation.

**Figure 6.**
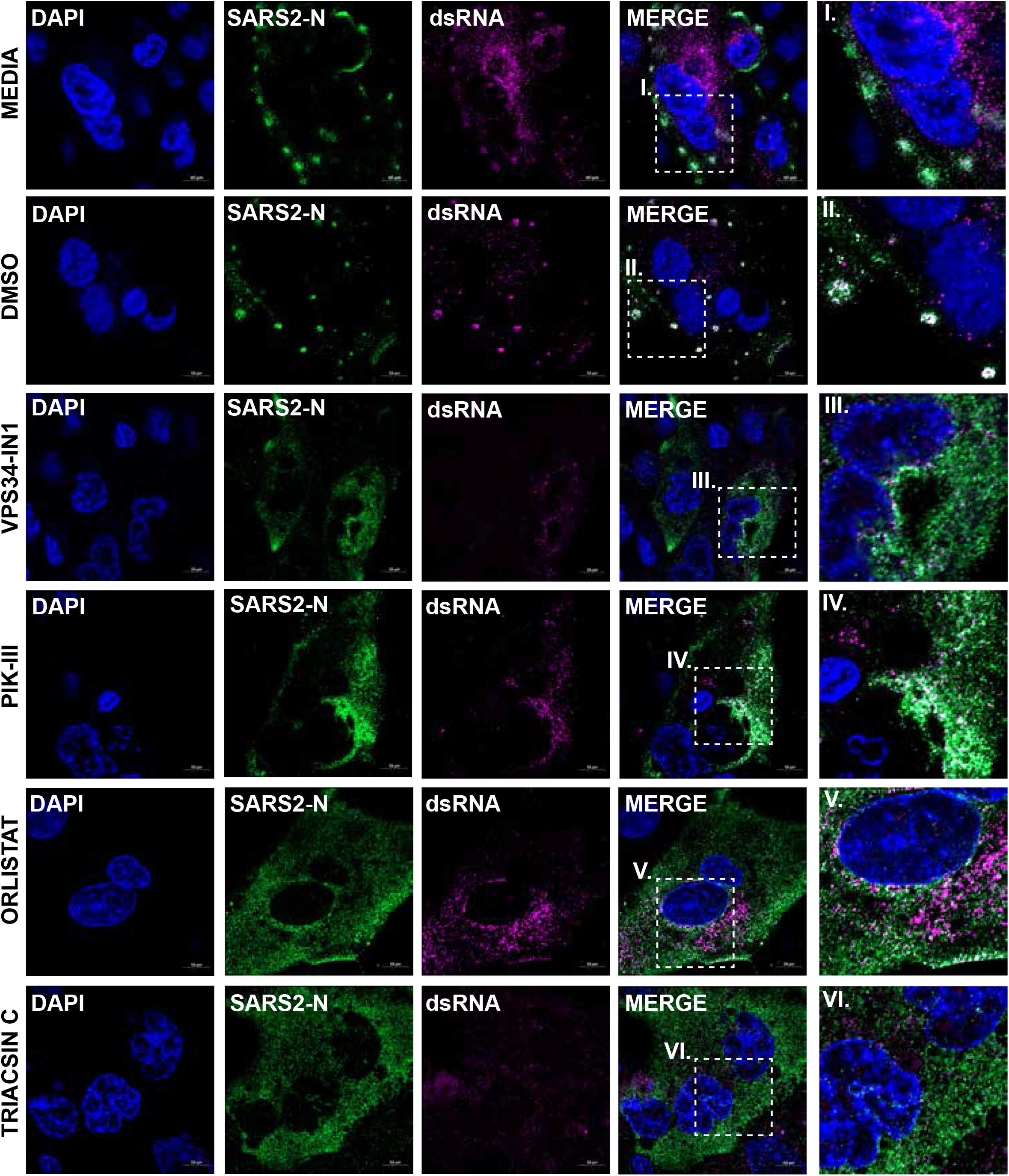
VPS34 activity and fatty acid metabolism are required to form SARS-CoV-2 N replication centers. Calu-3 cells were pre-treated with VPS34-IN1 (5uM), PIK-III (5uM), Orlistat (500uM), or Triacsin C (50uM) for 1 hour and infected with SARS-CoV-2 at MOI of 0.01. Cells were fixed at 24 h.p.i. and immunofluorescence was performed using primary antibodies against SARS-CoV-2 N or dsRNA, and AlexaFluor488 or AlexaFluor647 conjugated secondary antibodies, respectively. Nuclei were stained with Hoeschst 33342. Representative images are shown.

## DISCUSSION

Here, we demonstrate that two VPS34 inhibitors, Orlistat, and Triacsin C each have clear effects on SARS-CoV-2 replication and the morphology of viral replication centers. Generation of replication centers is a key feature of the replication of many viruses^36–38^. These can serve as sites where required components concentrate within a relatively closed environment and hide viral replication products from the host innate immune response^39^. In order to generate these centers, many viruses usurp host cellular pathways that are used to generate membranes or organelles^38^. *Betacoronaviruses* have been shown to target the ERAD-EDEMosome-ER pathways to generate double-membrane vesicles required for their replication^8^. The data presented here suggests roles for VPS34, FASN, and long chain fatty acyl CoA in replication center formation and stability suggesting a role for these host factors in providing the membranes needed for SARS-CoV-2 replication organelles.

VPS34 is of interest as a therapeutic target for a variety of conditions, including aging, neurodegeneration and cancer^40, 41^. The two VPS34 inhibitors tested were VPS34-IN1 and PIK-III which have *in vitro* IC50s for VPS34 of 25 nM and 18 nM, respectively^26, 42^. These were the most potent compounds versus SARS-CoV-2 tested in this study. Each displayed an IC50 of less than 1μM in either Vero E6 cells or Calu-3 cells. Activity in the Vero E6 cells was measured based on the capacity of the compounds to prevent viral cytopathic effects as measured by resistance across the cell monolayer, whereas the Calu-3 cell assay measured inhibition of production of infectious virus particles. The resistance-based assay provided a built-in measure of cell viability and integrity of the cell monolayer, providing assurance that decreases in resistance measurements initially post-infection were not reflective of cytopathic effects. We also independently determined that the compounds tested were non-toxic in Calu3 cells, likewise demonstrating that decreases in viral titer were not due to compound toxicity. Based on the Calu-3 data, the selectivity indices (SI) (CC50/IC50) for the compounds are >90 and >416 for VPS34-IN1 and PIK-III, respectively.

VPS34 is a phosphoinositide kinase that functions in autophagy, endosomal trafficking and other cellular functions^43^. VPS34 associates with VPS15 as well as with other proteins to carry out its activities. One VPS34-containing complex, Complex I, includes VPS34, VPS15, Beclin 1 and ATG14 and is critical for autophagosome formation. Complex II includes VPS34, VPS15, Beclin 1 and UVRAG and functions in autophagosome-lysosome fusion and in regulation of endosomes and multivesicular bodies^43^. While our inhibitor studies do not differentiate between the various functions of VPS34 that might be involved in SARS-CoV-2 replication,. Autophagy has been implicated as necessary for MHV replication, however, subsequent studies in different cell types suggest autophagy is not essential for MHV growth^7, 11^. Further, recent studies suggest that coronaviruses interfere with autophagy and that activation of autophagy can inhibit replication of SARS-CoV, MERS CoV, and SARS-CoV-2^44, 45^. Given that inhibition of VPS34 results in the inhibition of autophagy^26, 42^, it would be expected that inhibition of VPS34 would eliminate these anti-CoV effects of autophagy and promote SARS-CoV-2 replication. Therefore, the disruptions in SARS-CoV-2 replication due to VPS34 inhibition described here may, instead, reflect inhibition of non-autophagy related functions of VPS34.

Separate from autophagy, VPS34 has several other roles including in endosomal trafficking and retrograde endosome-to-Golgi transport^43^. For the positive-sense RNA virus TBSV, VPS34 was implicated in providing phosphatidylethanolamine-enriched membranes for formation of TBSV replication centers^13^. Based on our observation that VPS34 inhibitors disrupt the structure of SARS-CoV-2 replication centers, it is possible that VPS34 functions to facilitate membrane availability for SARS-CoV-2 replication organelle formation. Disruption of endocytic trafficking might also explain our observation that pre-treatment with VPS34 inhibitors alone had significant effects on SARS-CoV-2 replication.

Orlistat (tetrahydrolipstatin) is an FDA-approved weight loss drug that is taken orally and inhibits gastric and pancreatic lipases in the digestive tract, reducing uptake of lipids^21^. Orlistat also inhibits fatty acid synthase (FASN)^46^. Orlistat and other FASN inhibitors have previously been examined for their anti-cancer and antiviral activities. Although the clinically approved oral administration of Orlistat does not result in its significant systemic distribution, pre-clinical studies in mice have demonstrated that systemic administration of Orlistat is well tolerated^47^. Orlistat has been demonstrated to have activity against several viruses, including varicella-zoster virus (VZV), coxsackievirus B3 virus (CVB3), dengue virus (DENV), and other flaviviruses. DENV uses it nonstructural protein 3 to recruit FASN to viral replication sites and enhances synthesis of fatty acids^48^. As in our study, flaviviruses were sensitive to relatively high concentrations of Orlistat and antiviral effects could be demonstrated when Orlistat was added to cells post-infection^17^. Virus inhibition has typically been demonstrated at relatively high concentrations of Orlistat, such as 100μM or higher for CVB3, and between 10μM and 84μM for DENV3, depending on the timepoint post-infection DENV3 replication was measured^17, 19, 49^. For DENV3, the effect of Orlistat appeared to be after the early stages of infection^18^. This may reflect the need for DENV to recruit FASN to sites of virus replication and to upregulate fatty acid synthesis^48, 50^. It will be of interest to determine whether SARS-CoV-2 similarly depends on an upregulation of fatty acid synthesis.

Triacsin C inhibits long chain fatty acid acyl-CoA synthetase. Interestingly, the long chain fatty acid acyl-CoA synthetase ACSL3 was identified as an interactor of SARS-CoV-2 non-structural protein 7, suggesting a role for this enzyme in virus replication^51^. Triacsin C also has demonstrated antiviral activity for HCV and rotavirus^14–16^. For both HCV and rotavirus, the antiviral effects of Triacsin C have been linked to reliance of these viruses on lipid droplets for their replication^14–16^. Lipid droplets are organelles that store neutral lipids of which triglycerides are a major component^52^. By inhibiting long chain fatty acyl CoA, Triacsin C blocks lipid droplet formation. That antiviral activity against HCV and rotavirus is connected to lipid droplet formation is supported by the fact that these viruses are sensitive to inhibition by the DGAT inhibitors, T863 and PF06424439. In contrast, the compounds did not exhibit any activity against SARS-CoV-2 in Vero E6 cells whereas Triacsin C did. This suggests an alternate role for long chain fatty acyl CoA or its downstream metabolites other than triacylglycerol and lipid droplets. It is notable that the IC50 for Triacsin C was substantially lower in the Calu-3 cell assay as compared to the Vero cell assay. A lesser decrease in IC50 was also noted for Orlistat in the Calu-3 cells versus the Vero E6 cells. These observations may reflect different degrees of dependence of the virus on fatty acid metabolism in different cell types. From the perspective of antiviral development, it is encouraging that the human airway-derived cells are the more sensitive system given that SARS-CoV-2 targets the respiratory tract. Triacsin C has been administered to mice daily for up to two months without overt signs of significant toxicity and resulted in a decrease in atherosclerosis^53^. However, the pharmacokinetics and cell penetrance of Triacsin C are viewed as significant impediments to its clinical use^54^. Despite this, Triacsin C analogs have been developed^15^, and long chain fatty acyl CoA synthetases are of interest as potential therapeutics for cancer as well as for viruses^54^.

Cumulatively, these data support lipid metabolism as a potential therapeutic target for SARS-CoV-2 infection. The specific mechanisms by which VPS34 promotes SARS-CoV-2 replication and the precise manner in which the VSP34 inhibitors impair replication warrant further investigation. Additionally, the specific enzymes and products of fatty acid metabolism necessary for efficient SARS-CoV-2 growth in human airway epithelial cells should be further explored to more precisely identify relevant targets for therapeutic targeting. Further, it will be of interest to understand the relative efficacies of inhibitors of fatty acid metabolism in different cell types.

## Acknowledgments

This work was supported by NIH grants R01AI125453 and P01AI120943 (Amarasinghe) to CFB. We would like to thank the Georgia State University High Containment team Natasha Griffith, Martin Wildes, and Robert “Mike” Walsh for their continuous support.

---

**Supplemental Figure 1.**
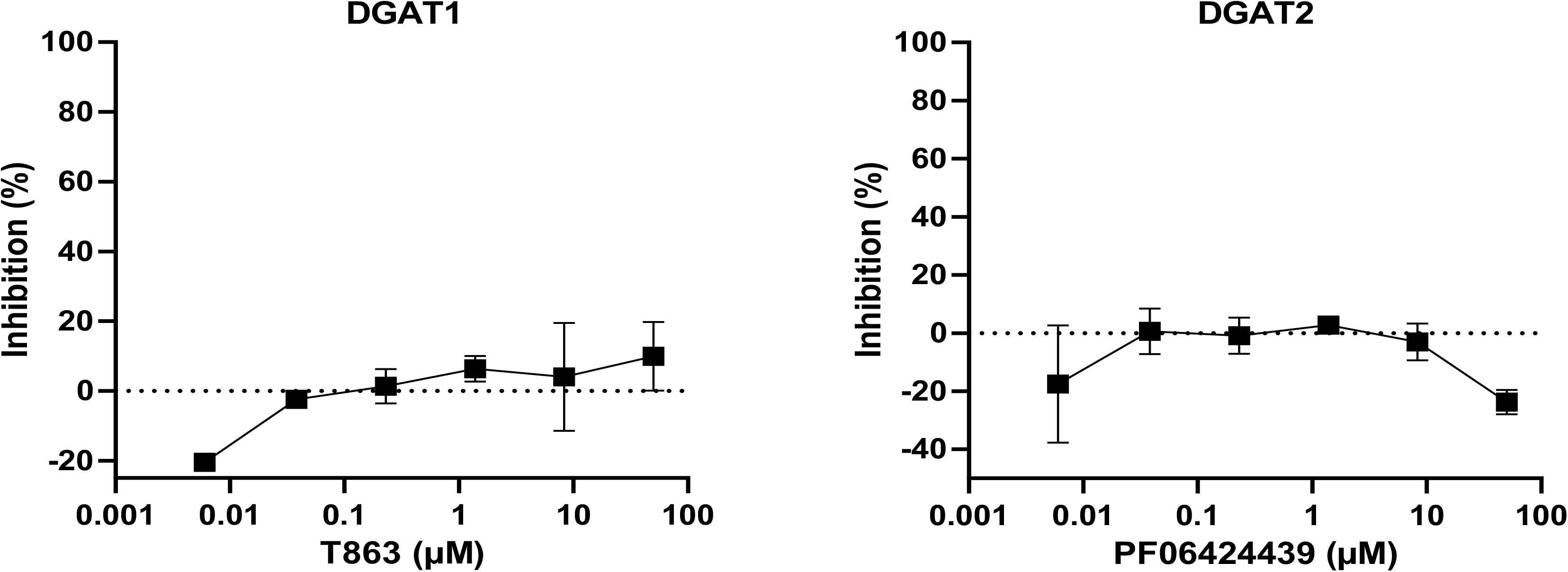
Inhibition of DGATs does not prevent SARS-CoV-2 replication. VeroE6 cells were seeded into a CytoView-Z 96-well plate and allowed to stabilize overnight. Cells were pre-treated with serial half-log dilutions of **A)** TC863 or **B)** PF06424439 and infected with SARS-CoV-2 at an MOI=0.01. Resistance was measured every minute over the course of 48 hours and percent inhibition relative to the DMSO control was determined at the 48-hour timepoint.

**Supplemental Figure 2.**
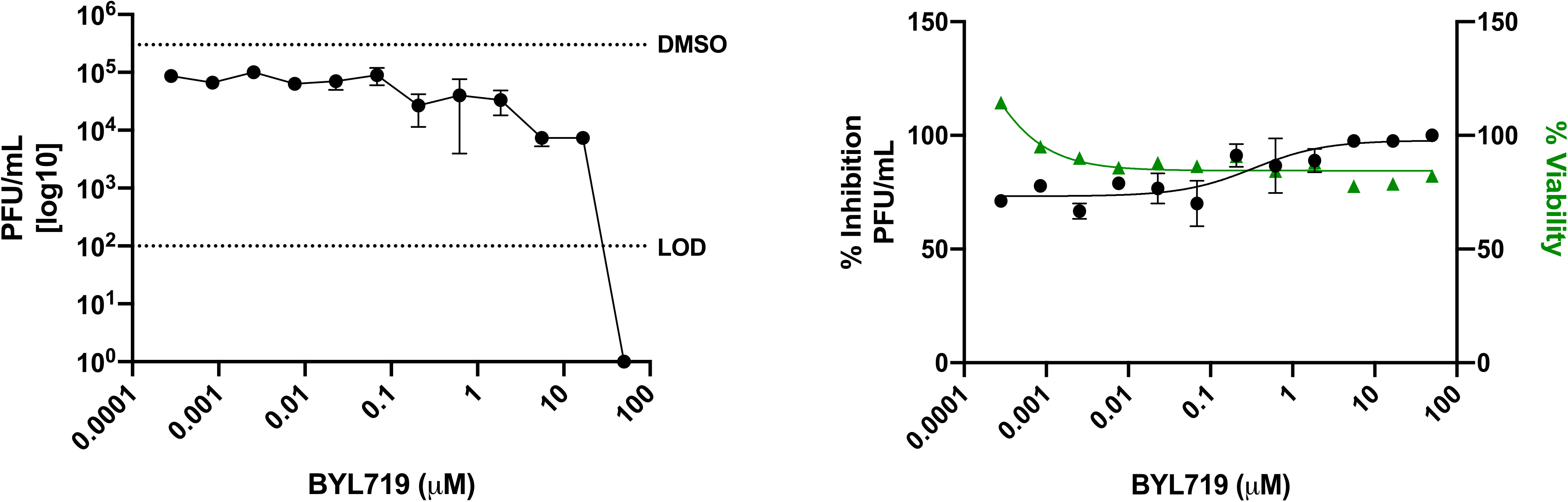
Inhibition of alpha PI3K does not prevent SARS-CoV-2 replication. Calu-3 cells were plated onto a 96-microplate and allowed to reach 95% confluency. Cells were then pre-treated with a range of concentrations of BYL719 and infected with SARS-CoV-2 at an MOI of 0.01. Supernatants were collected at 48 h.p.i. and titered on VeroE6 cells (left panel). Cell toxicity was determined in parallel and percent inhibition extrapolated from plaque assay data (right panel).

**Supplemental Figure 3.**
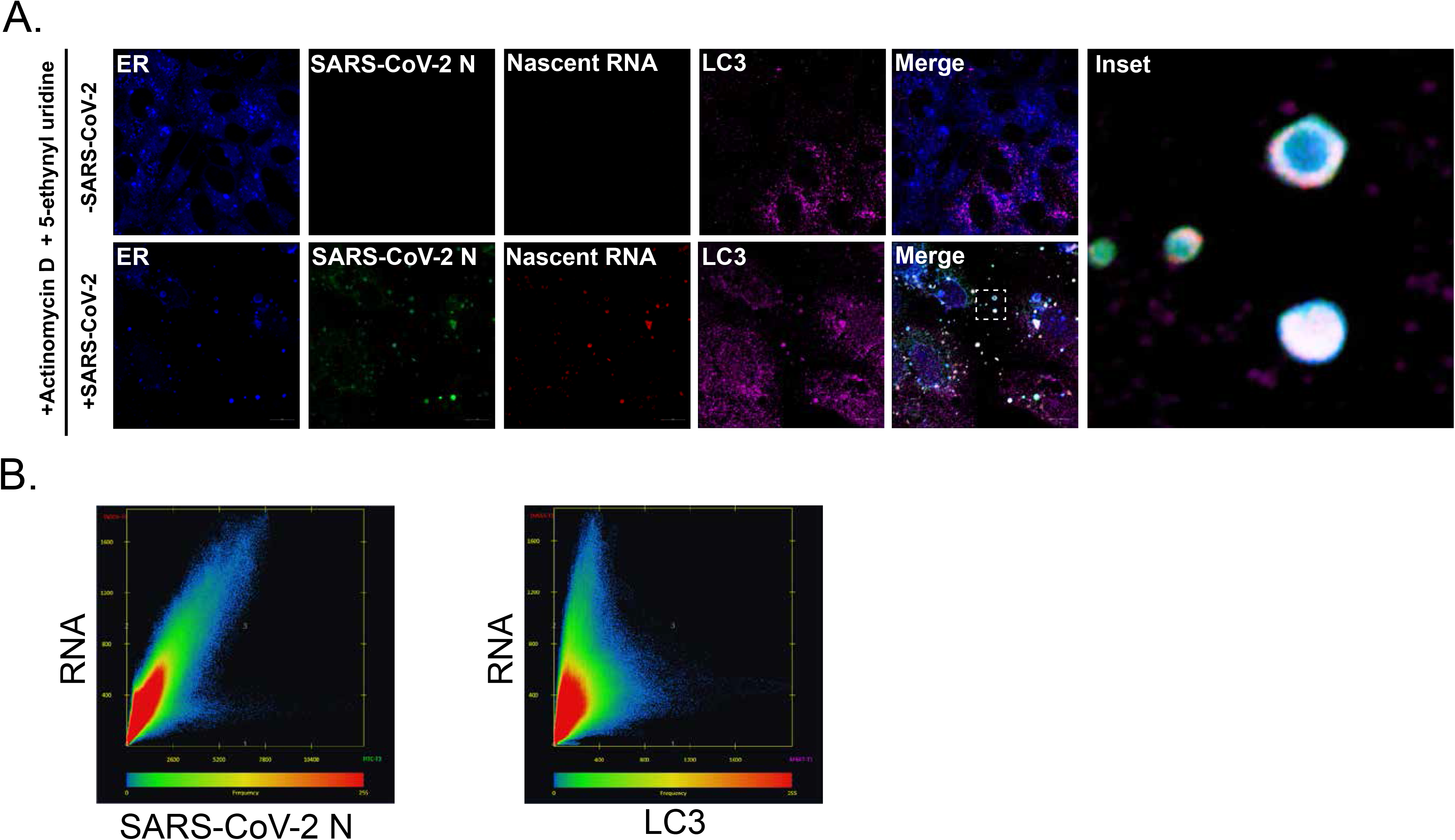
SARS-CoV-2 N and nascent viral RNA co-localize with the autophagy membrane marker LC3. VeroE6 cells were infected with SARS-CoV-2. At 24 h.p.i., cells were pre-treated with actinomycin D followed by a 5-ethynyl uridine (EU) chase for 4 hours. **A)** Cells were fixed, EU labeled viral nascent RNA was detected with click chemistry, and immunofluorescence performed using primary antibodies against SARS-CoV-2 N or LC3 and AlexaFluor488- or AlexaFluor647-conjugated secondary antibodies, respectively. Nuclei were stained with Hoeschst 33342. Representative images are shown. **B)** Co-localization was analyzed with Zen Blue.

## References

1. Lundstrom, K. Coronavirus Pandemic-Therapy and Vaccines. Biomedicines 8 (2020).

2. Wang, L., Wang, Y., Ye, D. & Liu, Q. Review of the 2019 novel coronavirus (SARS-CoV-2) based on current evidence. Int J Antimicrob Agents 55, 105948 (2020).

3. García-Serradilla, M., Risco, C. & Pacheco, B. Drug repurposing for new, efficient, broad spectrum antivirals. Virus Res 264, 22–31 (2019).

4. Pizzorno, A., Padey, B., Terrier, O. & Rosa-Calatrava, M. Drug Repurposing Approaches for the Treatment of Influenza Viral Infection: Reviving Old Drugs to Fight Against a Long-Lived Enemy. Front Immunol 10, 531 (2019).

5. Saini, K.S. et al. Repurposing anticancer drugs for COVID-19-induced inflammation, immune dysfunction, and coagulopathy. Br J Cancer (2020).

6. Li, G. & De Clercq, E. Therapeutic options for the 2019 novel coronavirus (2019-nCoV). Nat Rev Drug Discov 19, 149–150 (2020).

7. Prentice, E., Jerome, W.G., Yoshimori, T., Mizushima, N. & Denison, M.R. Coronavirus replication complex formation utilizes components of cellular autophagy. J Biol Chem 279, 10136–10141 (2004).

8. Reggiori, F. et al. Coronaviruses Hijack the LC3-I-positive EDEMosomes, ER-derived vesicles exporting short-lived ERAD regulators, for replication. Cell Host Microbe 7, 500–508 (2010).

9. Reggiori, F., de Haan, C.A. & Molinari, M. Unconventional use of LC3 by coronaviruses through the alleged subversion of the ERAD tuning pathway. Viruses 3, 1610–1623 (2011).

10. Snijder, E.J. et al. A unifying structural and functional model of the coronavirus replication organelle: Tracking down RNA synthesis. PLoS Biol 18, e3000715 (2020).

11. Zhao, Z. et al. Coronavirus replication does not require the autophagy gene ATG5. Autophagy 3, 581–585 (2007).

12. Su, W.C. et al. Rab5 and class III phosphoinositide 3-kinase Vps34 are involved in hepatitis C virus NS4B-induced autophagy. J Virol 85, 10561–10571 (2011).

13. Feng, Z., Xu, K., Kovalev, N. & Nagy, P.D. Recruitment of Vps34 PI3K and enrichment of PI3P phosphoinositide in the viral replication compartment is crucial for replication of a positive-strand RNA virus. PLoS Pathog 15, e1007530 (2019).

14. Liefhebber, J.M., Hague, C.V., Zhang, Q., Wakelam, M.J. & McLauchlan, J. Modulation of triglyceride and cholesterol ester synthesis impairs assembly of infectious hepatitis C virus. J Biol Chem 289, 21276–21288 (2014).

15. Kim, Y. et al. Novel triacsin C analogs as potential antivirals against rotavirus infections. Eur J Med Chem 50, 311–318 (2012).

16. Cheung, W. et al. Rotaviruses associate with cellular lipid droplet components to replicate in viroplasms, and compounds disrupting or blocking lipid droplets inhibit viroplasm formation and viral replication. J Virol 84, 6782–6798 (2010).

17. Hitakarun, A. et al. Evaluation of the antiviral activity of orlistat (tetrahydrolipstatin) against dengue virus, Japanese encephalitis virus, Zika virus and chikungunya virus. Sci Rep 10, 1499 (2020).

18. Tongluan, N. et al. Involvement of fatty acid synthase in dengue virus infection. Virol J 14, 28 (2017).

19. Ammer, E. et al. The anti-obesity drug orlistat reveals anti-viral activity. Med Microbiol Immunol 204, 635–645 (2015).

20. Esser, K. et al. Lipase inhibitor orlistat prevents hepatitis B virus infection by targeting an early step in the virus life cycle. Antiviral Res 151, 4–7 (2018).

21. Heck, A.M., Yanovski, J.A. & Calis, K.A. Orlistat, a new lipase inhibitor for the management of obesity. Pharmacotherapy 20, 270–279 (2000).

22. Benson, K., Cramer, S. & Galla, H.J. Impedance-based cell monitoring: barrier properties and beyond. Fluids Barriers CNS 10, 5 (2013).

23. Prentice, E., McAuliffe, J., Lu, X., Subbarao, K. & Denison, M.R. Identification and characterization of severe acute respiratory syndrome coronavirus replicase proteins. J Virol 78, 9977–9986 (2004).

24. Gordon, C.J. et al. Remdesivir is a direct-acting antiviral that inhibits RNA-dependent RNA polymerase from severe acute respiratory syndrome coronavirus 2 with high potency. J Biol Chem 295, 6785–6797 (2020).

25. Wu, J., Wu, B. & Lai, T. Compassionate Use of Remdesivir in Covid-19. N Engl J Med 382 (2020).

26. Bago, R. et al. Characterization of VPS34-IN1, a selective inhibitor of Vps34, reveals that the phosphatidylinositol 3-phosphate-binding SGK3 protein kinase is a downstream target of class III phosphoinositide 3-kinase. Biochem J 463, 413–427 (2014).

27. Wakil, S.J. & Abu-Elheiga, L.A. Fatty acid metabolism: target for metabolic syndrome. J Lipid Res 50 Suppl, S138–143 (2009).

28. Schütter, M., Giavalisco, P., Brodesser, S. & Graef, M. Local Fatty Acid Channeling into Phospholipid Synthesis Drives Phagophore Expansion during Autophagy. Cell 180, 135–149.e114 (2020).

29. Bakhache, W. et al. Fatty acid synthase and stearoyl-CoA desaturase-1 are conserved druggable cofactors of Old World Alphavirus genome replication. Antiviral Res 172, 104642 (2019).

30. Nasheri, N. et al. Modulation of fatty acid synthase enzyme activity and expression during hepatitis C virus replication. Chem Biol 20, 570–582 (2013).

31. Herker, E. et al. Efficient hepatitis C virus particle formation requires diacylglycerol acyltransferase-1. Nat Med 16, 1295–1298 (2010).

32. Cao, J. et al. Targeting Acyl-CoA:diacylglycerol acyltransferase 1 (DGAT1) with small molecule inhibitors for the treatment of metabolic diseases. J Biol Chem 286, 41838–41851 (2011).

33. Futatsugi, K. et al. Discovery and Optimization of Imidazopyridine-Based Inhibitors of Diacylglycerol Acyltransferase 2 (DGAT2). J Med Chem 58, 7173–7185 (2015).

34. Sims, A.C. et al. Severe acute respiratory syndrome coronavirus infection of human ciliated airway epithelia: role of ciliated cells in viral spread in the conducting airways of the lungs. J Virol 79, 15511–15524 (2005).

35. Sims, A.C., Burkett, S.E., Yount, B. & Pickles, R.J. SARS-CoV replication and pathogenesis in an in vitro model of the human conducting airway epithelium. Virus Res 133, 33–44 (2008).

36. Nagy, P.D., Strating, J.R. & van Kuppeveld, F.J. Building Viral Replication Organelles: Close Encounters of the Membrane Types. PLoS Pathog 12, e1005912 (2016).

37. Sasvari, Z. & Nagy, P.D. Making of viral replication organelles by remodeling interior membranes. Viruses 2, 2436–2442 (2010).

38. den Boon, J.A. & Ahlquist, P. Organelle-like membrane compartmentalization of positive-strand RNA virus replication factories. Annu Rev Microbiol 64, 241–256 (2010).

39. Santiago, F.W. et al. Hijacking of RIG-I signaling proteins into virus-induced cytoplasmic structures correlates with the inhibition of type I interferon responses. J Virol 88, 4572–4585 (2014).

40. Morris, D.H., Yip, C.K., Shi, Y., Chait, B.T. & Wang, Q.J. Beclin 1-Vps34 Complex Architecture: Understanding the Nuts and Bolts of Therapeutic Targets. Front Biol (Beijing) 10, 398–426 (2015).

41. Chude, C.I. & Amaravadi, R.K. Targeting Autophagy in Cancer: Update on Clinical Trials and Novel Inhibitors. Int J Mol Sci 18 (2017).

42. Dowdle, W.E. et al. Selective VPS34 inhibitor blocks autophagy and uncovers a role for NCOA4 in ferritin degradation and iron homeostasis in vivo. Nat Cell Biol 16, 1069–1079 (2014).

43. Backer, J.M. The intricate regulation and complex functions of the Class III phosphoinositide 3-kinase Vps34. Biochem J 473, 2251–2271 (2016).

44. Guo, L. et al. Autophagy Negatively Regulates Transmissible Gastroenteritis Virus Replication. Sci Rep 6, 23864 (2016).

45. Gassen, N.C. et al. SKP2 attenuates autophagy through Beclin1-ubiquitination and its inhibition reduces MERS-Coronavirus infection. Nat Commun 10, 5770 (2019).

46. Wakil, S.J. Fatty acid synthase, a proficient multifunctional enzyme. Biochemistry 28, 4523–4530 (1989).

47. Schcolnik-Cabrera, A. et al. Orlistat as a FASN inhibitor and multitargeted agent for cancer therapy. Expert Opin Investig Drugs 27, 475–489 (2018).

48. Heaton, N.S. et al. Dengue virus nonstructural protein 3 redistributes fatty acid synthase to sites of viral replication and increases cellular fatty acid synthesis. Proc Natl Acad Sci U S A 107, 17345–17350 (2010).

49. Wilsky, S. et al. Inhibition of fatty acid synthase by amentoflavone reduces coxsackievirus B3 replication. Arch Virol 157, 259–269 (2012).

50. Tang, W.C., Lin, R.J., Liao, C.L. & Lin, Y.L. Rab18 facilitates dengue virus infection by targeting fatty acid synthase to sites of viral replication. J Virol 88, 6793–6804 (2014).

51. Gordon, D.E. et al. A SARS-CoV-2 protein interaction map reveals targets for drug repurposing. Nature 583, 459–468 (2020).

52. Olzmann, J.A. & Carvalho, P. Dynamics and functions of lipid droplets. Nat Rev Mol Cell Biol 20, 137–155 (2019).

53. Matsuda, D. et al. Anti-atherosclerotic activity of triacsin C, an acyl-CoA synthetase inhibitor. J Antibiot (Tokyo) 61, 318–321 (2008).

54. Rossi Sebastiano, M. & Konstantinidou, G. Targeting Long Chain Acyl-CoA Synthetases for Cancer Therapy. Int J Mol Sci 20 (2019).

